# Generation of SARS-CoV-2 S1 spike glycoprotein putative antigenic epitopes in vitro by intracellular aminopeptidases

**DOI:** 10.1101/2020.06.22.164681

**Authors:** George Stamatakis, Martina Samiotaki, Anastasia Mpakali, George Panayotou, Efstratios Stratikos

**Affiliations:** Biomedical Sciences Research Center “Alexander Fleming”, Vari, Attica, Greece; National Centre for Scientific Research “Demokritos”, Agia Paraskevi, Attica, Greece

**Keywords:** Enzyme, Antigen, Peptide, Immune system, Aminopeptidase, Epitope, SARS-CoV-2, COVID-19, HLA, LC-MS/MS

## Abstract

Presentation of antigenic peptides by MHCI is central to cellular immune responses against viral pathogens. While adaptive immune responses versus SARS-CoV-2 can be of critical importance to both recovery and vaccine efficacy, how protein antigens from this pathogen are processed to generate antigenic peptides is largely unknown. Here, we analyzed the proteolytic processing of overlapping precursor peptides spanning the entire sequence of the S1 spike glycoprotein of SARS-CoV-2, by three key enzymes that generate antigenic peptides, aminopeptidases ERAP1, ERAP2 and IRAP. All enzymes generated shorter peptides with sequences suitable for binding onto HLA alleles, but with distinct specificity fingerprints. ERAP1 was the most efficient in generating peptides 8-11 residues long, the optimal length for HLA binding, while IRAP was the least efficient. The combination of ERAP1 with ERAP2 greatly limited the variability of peptide sequences produced. Less than 7% of computationally predicted epitopes were found to be produced experimentally, suggesting that aminopeptidase processing may constitute a significant filter to epitope presentation. These experimentally generated putative epitopes could be prioritized for SARS-CoV-2 immunogenicity studies and vaccine design. We furthermore propose that this *in vitro* trimming approach could constitute a general filtering method to enhance the prediction robustness for viral antigenic epitopes.

## Introduction

Severe acute respiratory syndrome coronavirus 2 (SARS-CoV-2) is the pathogen responsible for coronavirus disease 19 (COVID-19) that is behind a major ongoing pandemic (1–3). Virus entry into host cells is dependent on the S1 spike glycoprotein that forms homotrimers on the surface of the virion and interacts with the ACE2 receptor in susceptible cells (4–6).

Many studies on clinical characteristics and mortality resulting from SARS-CoV-2 infection have highlighted the need for a detailed understanding of immune responses against this pathogen (7). While appropriate innate and adaptive immune responses are necessary for recovery from infection, aberrant immune responses can be a major contributing factor to mortality (8, 9). In parallel, understanding immune recognition of SARS-CoV-2 is crucial to the ongoing massive global effort into developing an effecting vaccine against this pathogen (10). Although early analyses have focused on the development of neutralizing antibodies, cellular immune responses are emerging of vital importance (11) both for understanding normal immune response against this pathogen and for designing and optimizing vaccines (12). In particular T-cell mediated immunity appears to be important for both viral clearance and for long-term immunity (11). Thus, analysis of antigenic epitopes from SARS-CoV-2 should be a priority for the design of vaccines that induce effective and long-lasting cellular immune responses (13).

Cytotoxic T-cell responses against virus-infected cells hinge on the presentation of small peptidic fragments of viral proteins, called antigenic peptides, by specialized proteins on the cell surface that belong to the Major Histocompatibility Class I complex (MHCI, also called Human Leukocyte Antigens, HLA, in humans). Antigenic peptides are derived from viral proteins that are proteolytically degraded by complex proteolytic cascades (14). Intracellular aminopeptidases, ER aminopeptidase 1 (ERAP1), ER aminopeptidase 2 (ERAP2) and insulin-regulated aminopeptidase (IRAP) play important roles in producing antigenic peptides, by down-sizing longer peptides to the correct length for binding onto MHCI (15). Appropriate processing of pathogen antigens by these enzymes can determine the generation of cytotoxic immune responses and aberrant processing can lead to immune evasion (16). Thus, it is important to understand how these enzymes process SARS-CoV-2 antigens, so as to gain insight into the efficacy of antiviral cytotoxic responses and reveal possible avenues to enhance them.

In this study, we utilized a novel approach to analyze antigen trimming by intracellular aminopeptidases ERAP1, ERAP2 and IRAP, focusing on the largest antigen of SARS-CoV-2, namely S1 spike glycoprotein. By using tandem LC-MS/MS analysis, we were able to follow trimming in parallel of a large ensemble of peptides derived from the full length of S1 protein. This approach was inspired by two established observations: i) that these enzymes are expected to normally encounter a very large number of potential substrates concurrently in the cell and ii) accommodation of peptides inside a large cavity of each enzyme can lead to complex interactions between substrates that have to compete for the same space in the cavity (17–19). Our analysis provides novel insight into the differences in specificity between the three enzymes and provides a potential filter of traditional bioinformatic approaches that aim to predict antigenic epitopes. Finally, we propose a limited list of peptides that are potential ligands for common HLA alleles and could be prioritized for further immunological analyses and vaccine design efforts.

## Experimental procedures

### Enzyme expression and purification

Recombinant ERAP1, ERAP2 and IRAP were expressed and purified as described previously. Briefly, ERAP1 and ERAP2 were expressed by Hi5 insect cells in culture after infection with baculovirus carrying the appropriate gene and purified by affinity chromatography using a C-terminal his tag (20, 21). The enzymatic extracellular domain of IRAP was expressed by stably transfected HEK 293S GnTI^(−)^ cells and purified by affinity chromatography using a C-terminal Rhodopsin 1D4 tag (22). Enzymes were stored with 10% glycerol in aliquots at −80°C until needed.

### Peptides

The PepMix SARS-CoV-2 peptide mixture was purchased by JBT Peptide Technologies GmbH. Peptide pools were dissolved in DMSO. Prior to reactions the two peptide collections (158 and 157 peptides respectively) were mixed at equimolar concentrations and diluted in buffer containing 10mM Hepes pH 7, 100mM NaCl to a final concentration of 48μM.

### Enzymatic reactions

Enzymatic reactions were performed in triplicate in a total volume of 50μL in 10mM Hepes pH 7, 100mM NaCl. Freshly thawed enzyme stocks were added to each reaction to a final concentration of 50nM. Reactions were incubated at 37°C for 2 hours, stopped by the addition of 7.5μL of a 10% TFA solution, flash frozen in liquid nitrogen and stored at −80°C until analysis.

### LC-MS/MS analysis

The sample was preconcentrated on a pepmap LC trapping column (0.3X5mm) at a rate of 30uL of Buffer A (0.1% Formic acid in water) in 5min. The LC gradient used was 5% Buffer B (0.1% Formic acid in Acetonitrile) to 25% in 36 min followed by an increase to 36% in 5 min and a second increase to 80% in 0.5min and then was kept constant for 2min. The column was equilibrated for 15 min prior to the next injection. A full MS was acquired with a Q Exactive HF-X Hybrid Quadropole-Orbitrap mass spectrometer, in the scan range of 350-1400m/z using 60K resolving power with an AGC of 3× 10^6^ and max IT of 45ms, followed by MS/MS scans of the 12 most abundant ions, using 15K resolving power with an AGC of 1× 10^5^ and max IT of 22ms and an NCE of 28. The mass spectrometry proteomics data have been deposited to the ProteomeXchange Consortium via the PRIDE (23) partner repository with the dataset identifier PXD019901.

### Database Search

We employed the MaxQuant computational proteomics platform version 1.6.14.0 to search the peak lists against the Spike glycoprotein SARS2 FASTA file (Swissprot accession number P0DTC2) and a file containing 247 frequently observed contaminants. N-terminal acetylation (42.010565 Da) and methionine oxidation (15.994915 Da) were set as variable modifications. The second peptide identification option in Andromeda was enabled. The false discovery rate (FDR) were set to of 0.01 both for peptide. The enzyme specificity was set as unspecific. The minimum peptide length was set to 6 amino acids. The initial allowed mass deviation of the precursor ion was set to 4.5 ppm and the maximum fragment mass deviation was set to 20 ppm.

## Results and discussion

To investigate the trimming of antigenic epitope precursors by intracellular aminopeptidases that generate antigenic peptides, we used a mixture of 315 synthetic peptides derived from the sequence of the SARS-CoV-2 S1 spike glycoprotein. All peptides were 15 residues long and spanned the entire sequence of the protein with an 11 residue overlap between adjacent peptides. This mixture allows the systematic sampling of the entire sequence of the protein. The peptide mixture was incubated with either ERAP1, ERAP2, IRAP or an equimolar mixture of ERAP1 and ERAP2 at a substrate to enzyme ratio of 1000:1 and the digestion products analyzed by LC-MS/MS using a custom search database generated by *in silico* digestions of the full S1 protein sequence (UniProt ID: P0DTC2). All analyses were performed on three biological replicates for each reaction, as well as a control reaction that was performed in the absence of enzyme. An additional technical replicate for the control sample was also analyzed. For statistical robustness, we performed a t-test between the control sample and each reaction and selected for further analysis only the peptides for which the quantification value changed by a statistically significant degree (p-value<0.05).

The relative abundance of each peptide before and after the reaction was compared by label-free-quantification. Analysis identified 263 unique 15mers in the samples out of the total 315 in the mixture. This represents an 83% coverage of included peptides which may be due to poor ionization and detection for some peptide sequences. As a result, further analysis was limited to the peptides detectable by our experimental setup. On average, incubation with enzyme reduced the relative abundance of the 15mer peptides indicating successful digestion (Figure 1A, B). This reduction was much more evident for ERAP1 and ERAP2 (and their mixture) than for IRAP. Each enzyme featured a unique digestion fingerprint, suggesting different selectivity, as suggested in previous studies (24). Since the majority of peptides presented by HLA are 8-11 residues long, we analyzed the comparative abundance of 8-11mers generated from each reaction (Figure 1C, D). Of the three enzymes, ERAP1 was the most efficient in generating peptides within this length range, consistent with previous reports on the mechanism of action of this enzyme (17, 25). ERAP2 and the ERAP1/ERAP2 mixture followed, while IRAP was the least efficient. Similar to the trimming of 15mers, the generation of 8-11mers followed a unique fingerprint for each enzyme. This is consistent with the previous hypotheses that each of these enzymes accommodate peptides in a large internal cavity and selectivity is driven by interactions with the whole sequence of the peptide (17, 21, 22).

**Figure 1:**
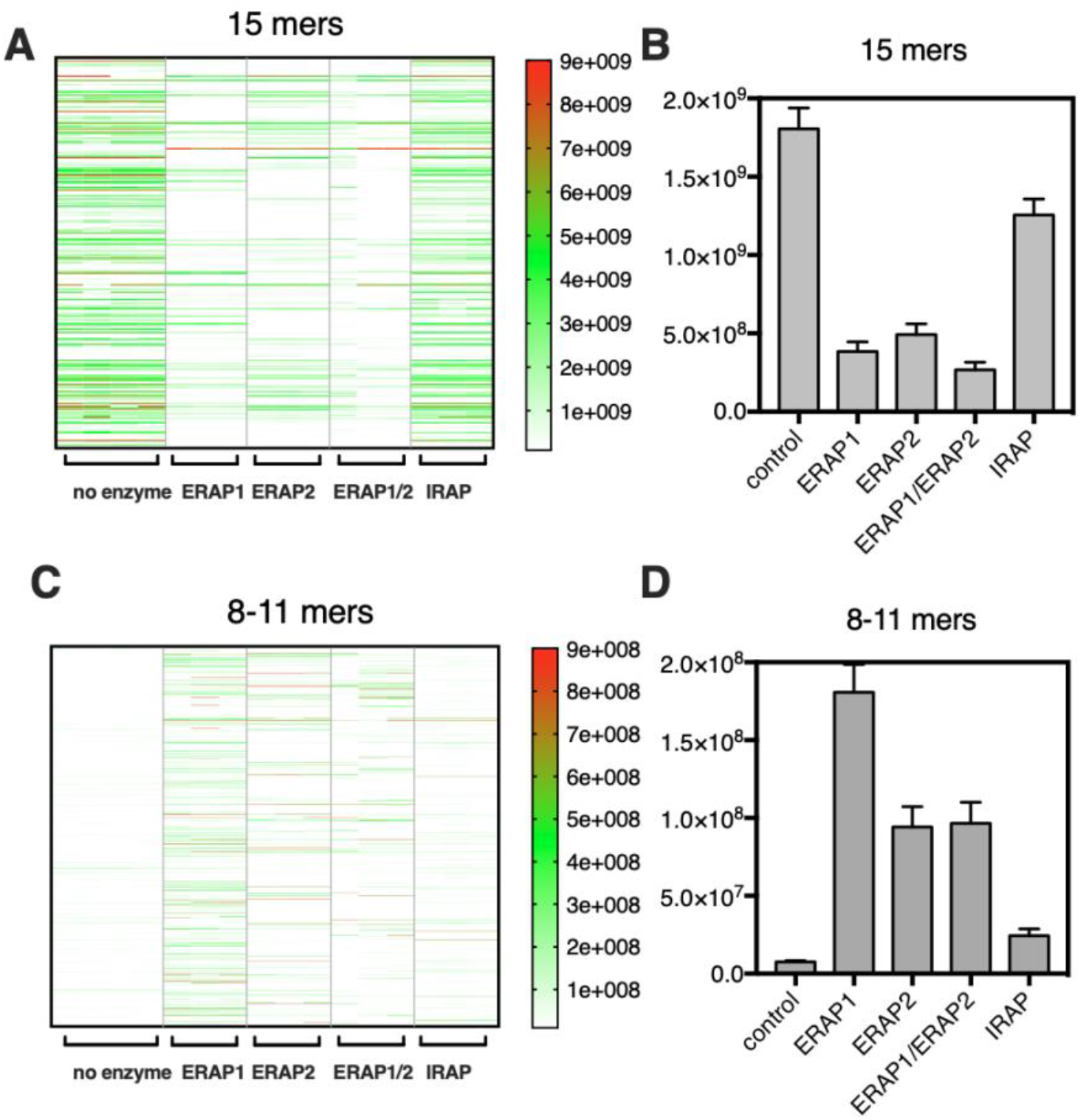
**Panel A**, heatmap showing trimming of 15mers by aminopeptidases for each biological replicate. **Panel B**, average label-free quantification (LFQ) signal of 15mers in the sample before and after incubation with the indicated enzyme. **Panel C**, heatmap showing LFQ signal for 8-11mers produced after digestion. **Panel D**, average signal of 8-11mers.

Indeed, comparing the peptide sequences generated by each enzyme, out of 1184 peptides identified, 142 were common between all three enzymes, 244 between ERAP1 and ERAP2 and 220 between ERAP1 and IRAP (Figure 2A). Furthermore, 169 peptides were unique for ERAP1, 234 for ERAP2 and 303 for IRAP. A similar situation was evident for 8-11mer peptides (Figure 2B). Strikingly, the mixture of ERAP1 with ERAP2 generated the fewest number of distinct sequences of 8-11mers (Figure 2C). This was in contrast to the finding that the ERAP1/ERAP2 mixture generated about the same average signal intensity as ERAP2 (Figure 1D). This was due to ERAP1/ERAP2 mixture generating fewer, in terms of sequence, distinct peptides, which were however relatively abundant. This finding is consistent with the proposed synergism of ERAP1 and ERAP2 (26) and suggests that the combination of these two enzymes is more efficient in trimming variable sequences and can thus over-trim peptides to lengths below 7 residues that are not detectable in our experimental setup and should not be able to stably bind onto MHCI (Figure 2C). As a result, incubation with ERAP1/ERAP2 mixture, accumulates only peptides that are resistant to degradation by both enzymes.

**Figure 2:**
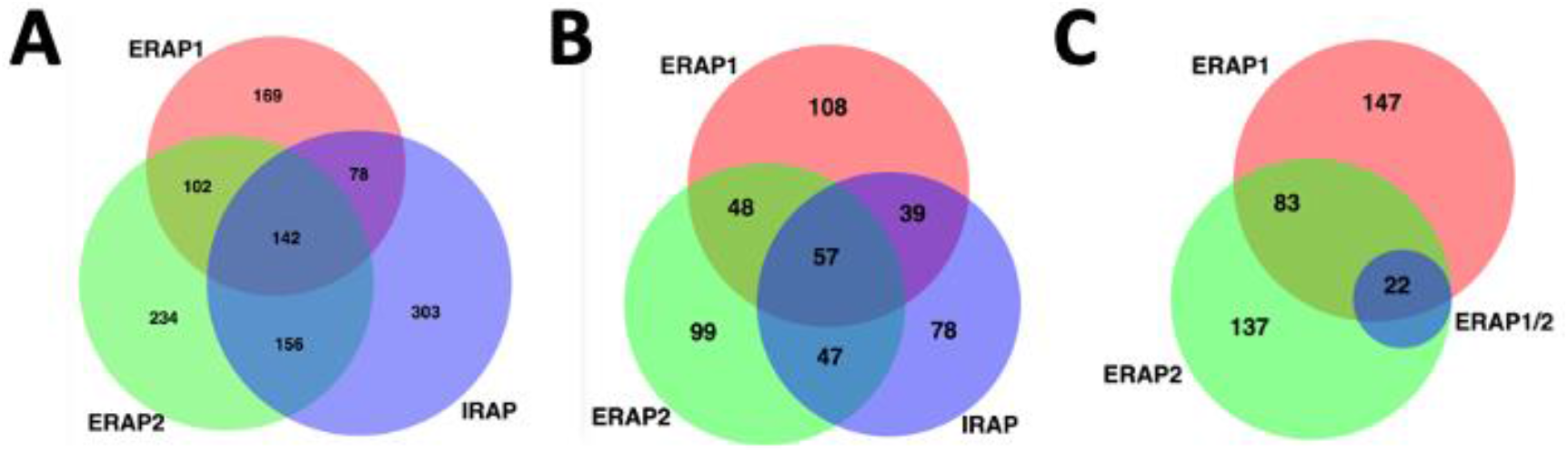
Venn diagrams indicating overlap between peptide sequences produced by each enzyme. Numerals indicate number of peptides in each segment. Analysis was performed using BioVenn (27). **Panel A**, comparison of all peptides produced by each enzyme. **Panel B,** comparison of 8-11mers produced by each enzyme. **Panel C**, comparison of 8-11mers produced by ERAP1, ERAP2 as well as their mixture (ERAP1/2).

Since epitope length is a key parameter for binding onto MHCI (the majority of presented peptides are 9mers) we analyzed the distribution of lengths of peptides generated by each enzymatic reaction (Figure 3). ERAP1 was very efficient in trimming the 15mer substrates and generated primarily 9mer peptides, consistent with its proposed property as a “molecular ruler” (25). Neither ERAP2 nor IRAP were able to accumulate 9mers preferably, but still generated significant numbers. The mixture of ERAP1 and ERAP2 showed a similar fractional distribution of peptide lengths (Figure 3B), but produced a much lower number of distinct peptide sequences (Figure 3A), presumably due to over-trimming to smaller lengths or even single amino acids.

**Figure 3:**
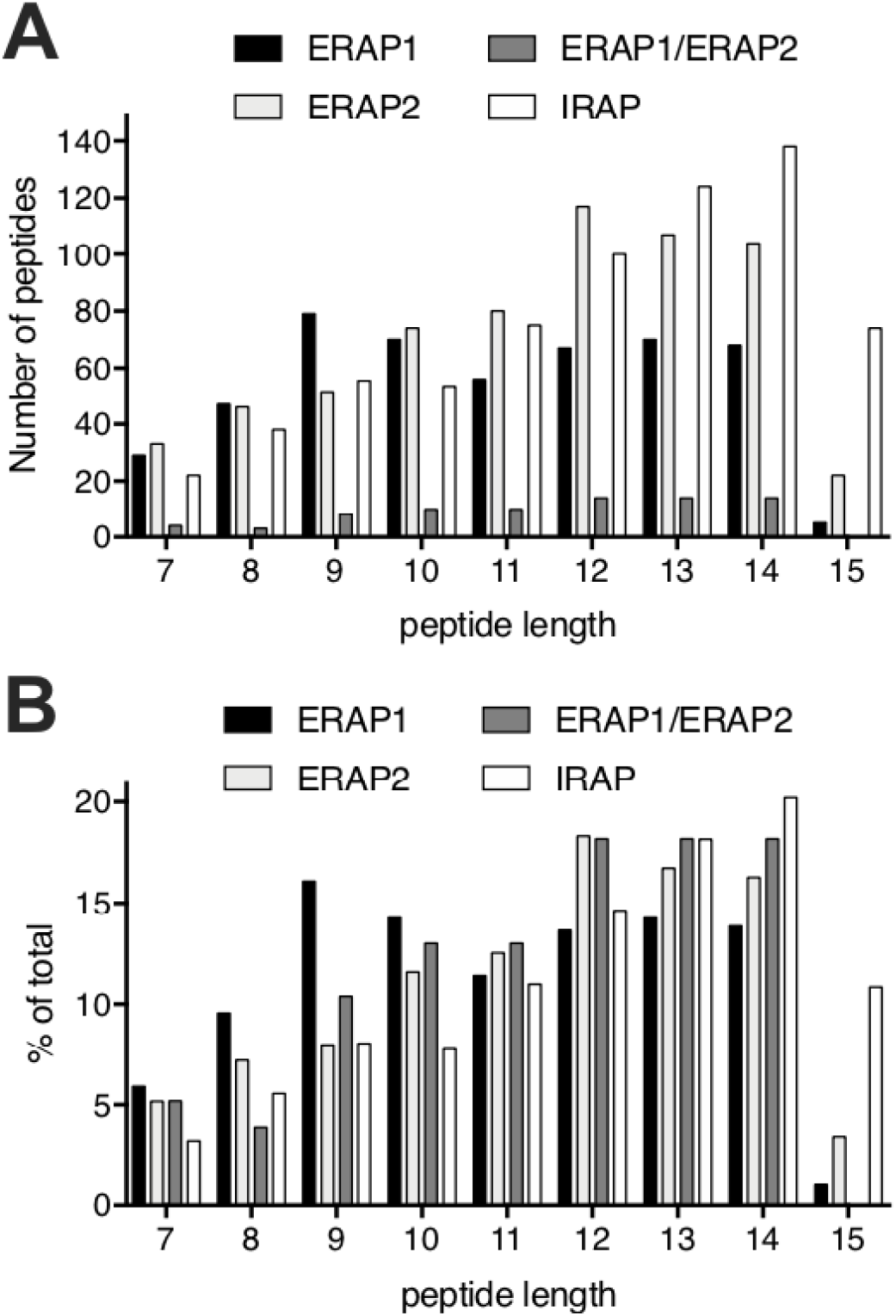
Length distribution of peptides detected by LC-MS/MS after enzymatic digestion. **Panel A,** number of peptides detected of each length. **Panel B,** relative fraction of each length compared to the total number of peptides detected.

The main determinant in antigen presentation is stable binding of antigen-generated peptides onto MHCI. To evaluate the potential of the generated peptides to bind onto MHCI we used the HLAthena prediction server to rank the peptides for binding onto a collection of common HLA alleles (28) specifically HLA-A01:01, HLA-A02:01, HLA-A03:01, HLA-A24:02, HLA-A26:01, HLA-B07:02, HLA-B08:01, HLA-B27:05, HLA-B39:01, HLA-B40:01, HLA-B58:01 and HLA-B15:01 (Supplemental Table 1). For each peptide we selected the best scoring HLA-allele and plotted the calculated percentile rank of the predicted score for each enzymatic reaction (Figure 4A). The geometric mean of the predicted affinity was lowest for ERAP1 (indicating that the ERAP1 generated peptides had the highest average affinity for HLA), followed by IRAP and then ERAP2. Only a subset of generated peptides was predicted to bind with sufficient affinity onto at least one HLA: 23% for ERAP1, 22% for ERAP2, 6% for ERAP1/ERAP2 mixture and 21% for IRAP (peptide sequences are listed in Table 1a). These peptides spanned the whole sequence of the S1 spike glycoprotein, although each enzyme presented a unique signature onto this sequence (Figure 4B).

**Figure 4:**
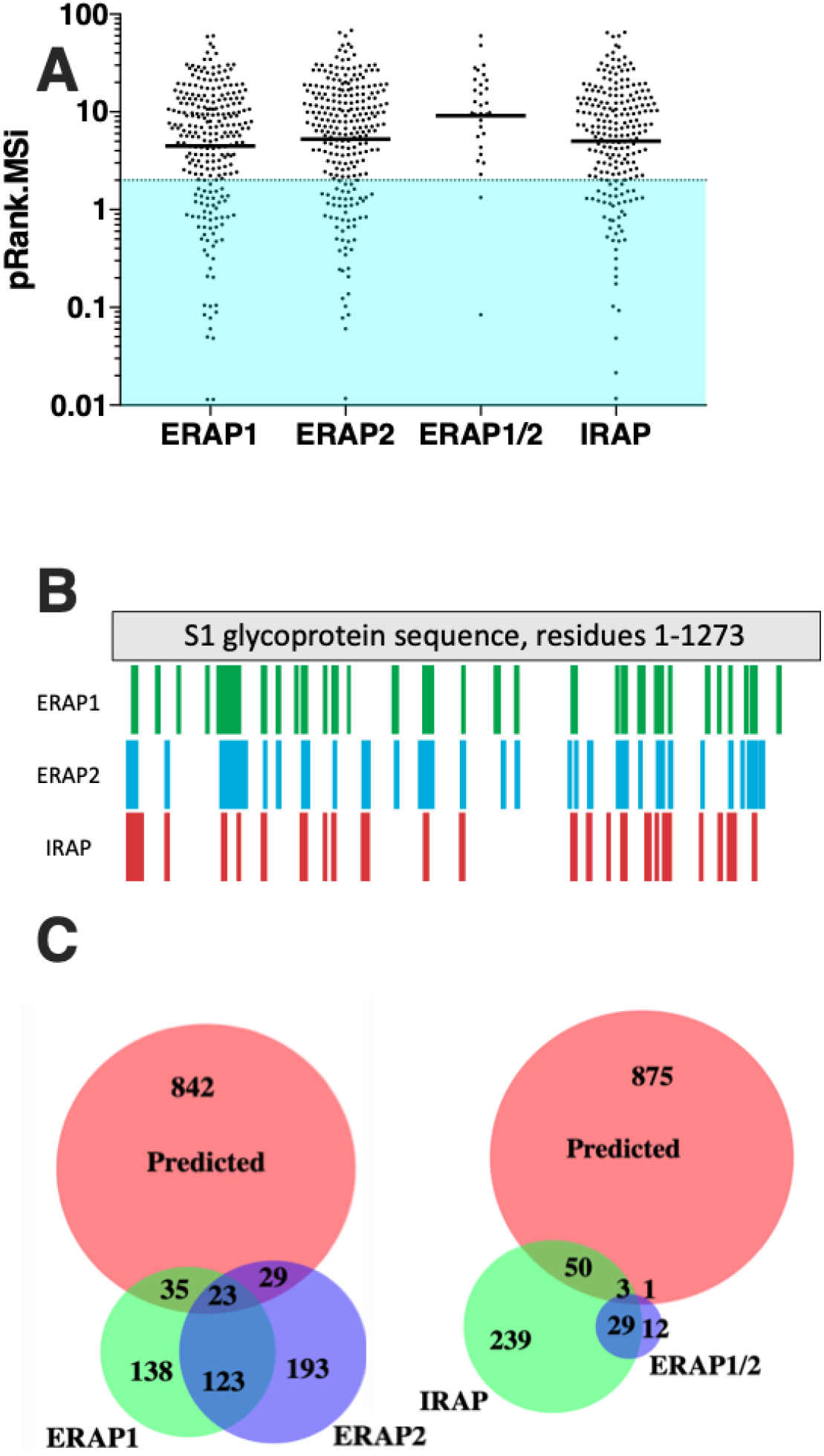
**Panel A,** scatter plot showing the predicted affinity of produced peptides for common HLA alleles as calculated by HLAthena. Color region encompasses peptides that are predicted to bind to at least one of the common HLA alleles used in the analysis. **Panel B,** schematic representation of relative locations in the S1 protein sequence where the generated peptides are found. **Panel C,** Venn diagrams depicting overlap between peptides of S1 protein predicted to bind to common HLA alleles and peptides produced experimentally by ERAP1, ERAP2, IRAP or ERAP1/ERAP2 mixture. Numerals indicate number of peptides in each separate segment.

**Table 1a:**
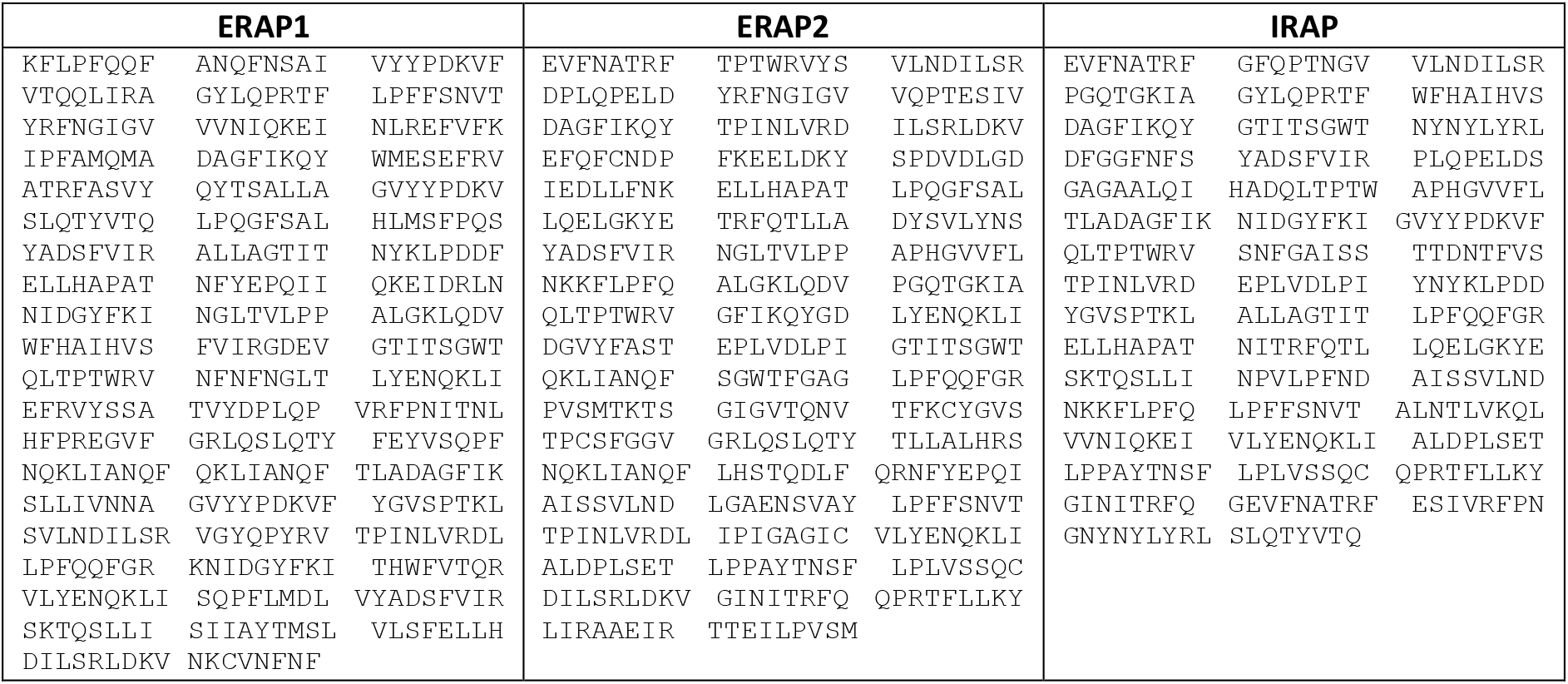
Peptides generated by each aminopeptidase and are predicted to bind onto common HLA-alleles.

In a recent publication, the authors proposed that different HLA alleles can have significant variability in their ability to present SARS-CoV-2 epitopes, with HLA-B46:01 having the capability to present the fewest and HLA-B15:03 being able to present the most (29). We thus used HLAthena to analyze the propensity of the experimentally produced peptides to bind onto these two alleles (Supplemental Table 2). ERAP1 was found to produce 15 potential ligands for HLA-B15:03 but only 6 for HLA-B46:01, consistent with the proposed trend. In contrast, ERAP2 produced 8 potential ligands for both alleles and IRAP produced 6 for HLA-B15:03 and 4 for HLA-B46:01. Strikingly, the mixture of ERAP1 with ERAP2 produced 4 peptides that could bind onto HLA-B15:03, but no peptides predicted to bind onto HLA-B46:01 (Table 1b). Thus, our findings appear to validate the hypothesis that HLA-B15:03 is likely to present more SARS-CoV-2 epitopes than HLA-B46:01, but only for ERAP1, which however is considered the dominant aminopeptidase activity in the cell for generating antigenic peptides.

**Table 1b:**
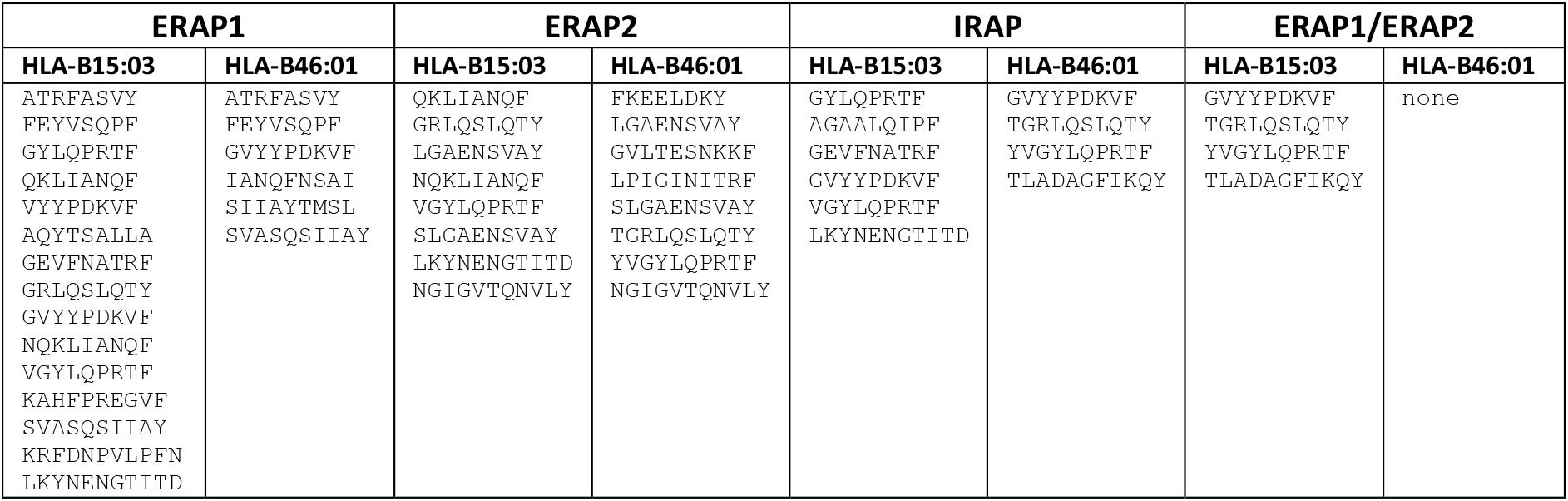
Peptides generated by each aminopeptidase that are predicted to bind to HLA-B15:03 and HLA-B46:01

| ERAP1 |  | ERAP2 |  | IRAP |  | ERAP1/ERAP2 |  |
| --- | --- | --- | --- | --- | --- | --- | --- |
| HLA-B15:03 | HLA-B46:01 | HLA-B15:03 | HLA-B46:01 | HLA-B15:03 | HLA-B46:01 | HLA-B15:03 | HLA-B46:01 |
| ATRFASVY | ATRFASVY | QKLIANQF | FKEELDKY | GYLQPRTF | GVYYPDKVF | GVYYPDKVF | none |
| FEYVSQPF | FEYVSQPF | GRLQSLQTY | LGAENSVAY | AGAALQIPF | TGRLQSLQTY | TGRLQSLQTY |  |
| GYLQPRTF | GVYYPDKVF | LGAENSVAY | GVLTESNKKF | GEVFNATRF | YVGYLQPRTF | YVGYLQPRTF |  |
| QKLIANQF | IANQFNSAI | NQKLIANQF | LPIGINITRF | GVYYPDKVF | TLADAGFIKQY | TLADAGFIKQY |  |
| VYYDPKVF | SIIAYTMSL | VGYLQPRTF | SLGAENSVAY | VGYLQPRTF |  |  |  |
| AQYTSALLA | SVASQSIIAY | SLGAENSVAY | TGRLQSLQTY | LKYNENGTITD |  |  |  |
| GEVFNATRF |  | LKYNENGTITD | YVGYLQPRTF |  |  |  |  |
| GRLQSLQTY |  | NGIGVTQNVLY | NGIGVTQNVLY |  |  |  |  |
| GVYYPDKVF |  |  |  |  |  |  |  |
| NQKLIANQF |  |  |  |  |  |  |  |
| VGYLQPRTF |  |  |  |  |  |  |  |
| KAHFPREGVF |  |  |  |  |  |  |  |
| SVASQSIIAY |  |  |  |  |  |  |  |
| KRFDNPVLPFN |  |  |  |  |  |  |  |
| LKYNENGTITD |  |  |  |  |  |  |  |

Bioinformatic epitope predictions based on antigen sequence are often used as a tool to study the potential antigenicity of a particular epitope or pathogen. The power of those predictions is constantly evolving and primarily relies on predictions of binding affinity on HLA. To compare such predictions to our experimentally generated peptides, we used the full sequence of the S1 spike glycoprotein and the NetMHCpan 4.1 server (30) to predict potential epitopes that could be presented by the common HLA alleles indicated in the previous paragraph. The server predicted 929 potential epitopes with lengths of 8-12 residues (Supplemental Table 3). Of those potential epitopes however, less than 7% were found to be produced experimentally by one of the enzymes tested and more specifically 58 by ERAP1, 52 by ERAP2, 4 for the ERAP1/2 mixture and 53 by IRAP (Figure 4C). While our experimental approach has limitations as discussed below, this finding suggests that intracellular antigen processing by aminopeptidases may constitute a major filter in determining which peptides will be presented by MHCI. Indeed, it has been recently proposed that the main function of ERAP1 is to limit the peptide pool available for MHCI (31). In this context, this experimental approach could be useful in optimizing bioinformatic predictions.

Our findings provide new information on both the general biological functions of intracellular aminopeptidases that generate antigenic peptides as well as on specific processing of a key antigen from SARS-CoV-2. Specifically, our results highlight that each enzyme bears a unique trimming fingerprint to antigen processing. Although this has been suggested before based on differences in specificity towards specific peptide substrates (26, 32–34), it has not been observed in the context of peptide ensembles. This is potentially important since competition of different peptides for the cavity in these enzymes could result in complex substrate interactions. At first glance, major differences in trimming fingerprints between each or the three enzymes, may appear to impose an unnecessary complication to antigenic peptide generation. It is conceivable, however, that this trimming variability is desirable for the immune system, since it can expand the breadth of possible antigenic peptides detected in different immunological settings and cell types. Our results also highlight a previously proposed property of ERAP1: the specialization in trimming large peptides and producing peptides that have the ideal length for MHCI binding – most of the ERAP1 products fall well within that range (25). In contrast, both ERAP2 and IRAP appear to be less optimized for length selection. However, they are still able to produce many peptides that are potential cargo for MHCI, casting some doubt on whether the unique trimming properties of ERAP1 are absolutely necessary for this basic function. Furthermore, the combination of ERAP1 with ERAP2 appears to provide significant synergism in trimming, to the point of over-trimming peptides and limiting available sequences. Synergism between ERAP1 and ERAP2 has been demonstrated before in trimming isolated peptides and these two enzymes have been proposed to also form functional dimers (33, 35). According to our observations, their combination is especially efficient in trimming. While the biological repercussions of this are not fully clear yet, it is conceivable that the strong associations between ERAP2 activity and predisposition to autoimmunity may be related to this effect (36).

Despite the current importance in understanding immune reactions in COVID-19, very little is known about the cellular adaptive immune responses against SARS-CoV-2. Cellular immune responses are emerging as a central player in clearing the infection and as targets for vaccine efforts (37, 38). Furthermore, HLA polymorphic variation has been suggested to underlie the large variability in virus clearance that has been observed amongst individuals (39). Our analysis of the largest antigen of SARS-CoV-2, S1 spike glycoprotein, suggests that aminopeptidase trimming can be a significant filter that helps shape which peptides will be presented by HLA. Thus, we propose a short list of candidate peptides that could be prioritized in downstream antigenic analysis as well as in vaccine design and efficacy studies.

While the functions of ERAP1, ERAP2 and IRAP have been studied in both *in vitro* and *in vivo* contexts during the last decade, their relative functional differences have only been compared in processing specific substrates at a time. However, all these enzymes have a broad substrate specificity and can normally encounter thousands of different peptides in the ER or endosomal compartments. On the other hand, studies focusing on the presented immunopeptidome have revealed effects attributed to ERAP1 and ERAP2 trimming, but direct comparisons have been difficult because of the dominant effect of MHCI affinity on presentation (40). Our approach stands in-between these two types of studies. It mimics the multiple-substrate situation that is likely normal *in vivo* but focuses on antigenic peptide precursor trimming. In this context, our approach may have broader application for the quick prediction of potential antigenic epitopes as an additional filter on bioinformatic predictions. Indeed, bioinformatic predictions result in many candidate peptides, very few of which will provoke an immune response; adding more filters can increase the usefulness of these rapid approaches. However, our approach also has limitations that need to be taken into account when interpreting results. Due to differences in ionization and detection by the LC-MS/MS some peptides may not be detected or may be under-represented compared to other sequences, making comparisons between different peptides less reliable. Furthermore, it is an *in vitro* approach that is limited to the peptide pool used and cannot take into account the dynamics of MHCI binding that can protect peptides from further aminopeptidase degradation (41) or peptide proofreading by chaperone components or the peptide loading complex (42). Due to those limitations, we restricted our analysis to statistical comparisons and avoided drawing conclusions regarding particular peptide sequences.

In summary, we analyzed the trimming of a peptide ensemble spanning the sequence of the S1 spike glycoprotein of SARS-CoV-2, the pathogen responsible for the recent COVID-19 pandemic. Our analysis provided novel insight into the function of antigen trimming enzymes and suggested that aminopeptidase trimming may be a significant filter in determining which peptides can be presented by MHCI. Furthermore, we have identified a limited set of peptides that were experimentally produced by elongated precursors which could be prioritized in future studies aiming to investigate the antigenicity of SARS-CoV-2 infected cells and assist in the design of highly effective vaccines that aim to produce adaptive cytotoxic responses. We propose that our experimental approach may also be useful as a general tool for enhancing bioinformatic predictions of antigenic epitopes.

## Notes

### Competing Interest Statement

The authors have declared no competing interest.

